# DMC-BrainMap - an open-source, end-to-end tool for multi-feature brain mapping across species

**DOI:** 10.1101/2025.02.19.639009

**Authors:** Felix Jung, Xiao Cao, Loran Heymans, Marie Carlén

## Abstract

A central goal of current neuroscience research is to understand how behavior emerges from neuronal circuit activity. For this, cellular and circuit components in intact brains need to be identified and studied using e.g. viral tracing, electrophysiology, imaging, activity perturbations, and behavior. A commonality of these approaches is the need to establish the anatomical locations of both experimental parameters (e.g. recording/injection sites) and circuit components (e.g. cell bodies/projections). The development of standardized coordinate systems, i.e. reference brain atlases, has proven essential for reproducible, cross-laboratory mapping of anatomical data. However, the process of mapping, analyzing, and visualizing anatomical data remains challenging, especially for users lacking programming expertise. Here, we introduce *DMC-BrainMap*, an open-source *napari* plugin designed as a user-friendly tool for streamlined processing and whole-brain analysis of anatomical data. The working principle and core functionalities include all steps after image acquisition, i.e., preprocessing of images, registration of images to a reference atlas, segmentation of different anatomical features, and data analysis/visualization. *DMC-BrainMap* can be applied to histological data obtained from a variety of model organisms at different developmental stages, including mouse, rat, and zebrafish. We demonstrate the utility of *DMC-BrainMap* by mapping and quantifying cell bodies, axonal densities, injection sites as well as placement of optical fibers and Neuropixels probes. Further, we show how spatial transcriptomics data can be integrated into the *DMC-BrainMap* workflow. By eliminating the need for programming by the user, *DMC-BrainMap* provides an easy-to-use tool for increased rigor, reproducibility, and data sharing in neuroscientific research involving animal models.

## Introduction

Understanding how computations underlying complex behaviors are implemented in brain-wide circuits is a central goal of current neuroscience research [1–3]. For this, viral tracing, large-scale *in vivo* electrophysiology and imaging, activity perturbations, and ‘spatial-omics’ have become standardized tools in neuroscience laboratories to study circuit components and their physiology. A commonality of current studies is the need for anatomical mapping of the data, e.g. the location of cell bodies, implanted electrodes and optical fibers, and the spread of viral vectors. A variety of tools for mapping anatomical data to standardized coordinate systems, so-called reference atlases, has been developed in recent years, tailored to specific anatomical features and tissue preparations (brain sections (2D data) versus volumetric (3D) data; Table in S1 Table). With a focus on volumetric data, the BrainGlobe Initiative (https://brainglobe.info/) has generated tools allowing users to segment [4] and register anatomical features to reference atlases [5] and create high-quality data visualizations ([6]; see also [7–10]). However, brain sections are still the most commonly used preparation [11]. Several tools are available for registration of images of (mouse) brain slices to a reference atlas and mapping of anatomical data, all holding their specific advantages and limitations [10–17]. Most tools, however, do not cover the whole process from image preprocessing to data visualization. Other common limitations are the need for multiple or proprietary software and programming skills. To overcome these hurdles, we developed *DMC-BrainMap*, a plugin for the Python-based image viewer *napari* (https://napari.org/). *DMC-BrainMap* is an interactive end-to-end tool tailored for 2D brain section data that covers all steps from image preprocessing to data visualization. We also provide *DMC-FluoImager,* a software for scanning 2D brain sections (https://github.com/hejDMC/dmc-fluoimager). The *DMC-BrainMap* pipeline is installed, run, and controlled through a graphical user interface (GUI), eliminating the need for programming by the user. The pipeline focuses on user-friendly operation and facilitated manual curation of image registration and anatomical feature segmentation. As the pipeline integrates the BrainGlobe Atlas API [18] it can be applied to data from e.g. mice, rats, and zebrafishes at different developmental stages. The provided pycro-manager-based [19] *DMC-FluoImager* software can be used for customized scanning of 2D brain sections using any microscope equipped with a motorized XYZ stage. Data obtained via *DMC-FluoImager* is natively structured for seamless use within the *DMC-BrainMap* workflow. Alternatively, any single channel 16-bit .tif/.tiff images can be used as input to the *DMC-BrainMap* pipeline. The *DMC-BrainMap* pipeline provides solutions for convenient batch preprocessing of the microscopic images, fast and interactive image registration to reference atlases, and semi-automatic segmentation of a variety of anatomical features, including (labeled) cell bodies, axonal densities, injection sites, and tracts from optical fibers or Neuropixels probes 1.0. The integrated software *ProbeViewer* offers voxel-wise inspection of reconstructed Neuropixels tracts. The pipeline in addition enables users to analyze and visualize data, including spatial transcriptomics (ST) data, in a variety of ways. The *DMC-BrainMap* pipeline is open-source, compatible with Windows, Mac, and Linux operating systems, and can be integrated with other *napari* plugins.

**Table S1. List of tools for mapping of anatomical data**.

## Results

The *DMC-BrainMap* pipeline is designed as a user-friendly tool for streamlined processing and whole-brain analysis of anatomical data. The working principle and core functionalities include all steps after image acquisition, i.e., preprocessing of images, registration of images to a reference atlas, segmentation of anatomical features, and data analysis/visualization (Fig 1A-B). We also provide a solution for image acquisition, *DMC-FluoImager*, a script for automatic imaging of 2D brain sections with a fluorescence microscope (Fig 1A). The output data format and structure of *DMC-FluoImager* are readily compatible with the *DMC-BrainMap* pipeline.

**Fig 1.**
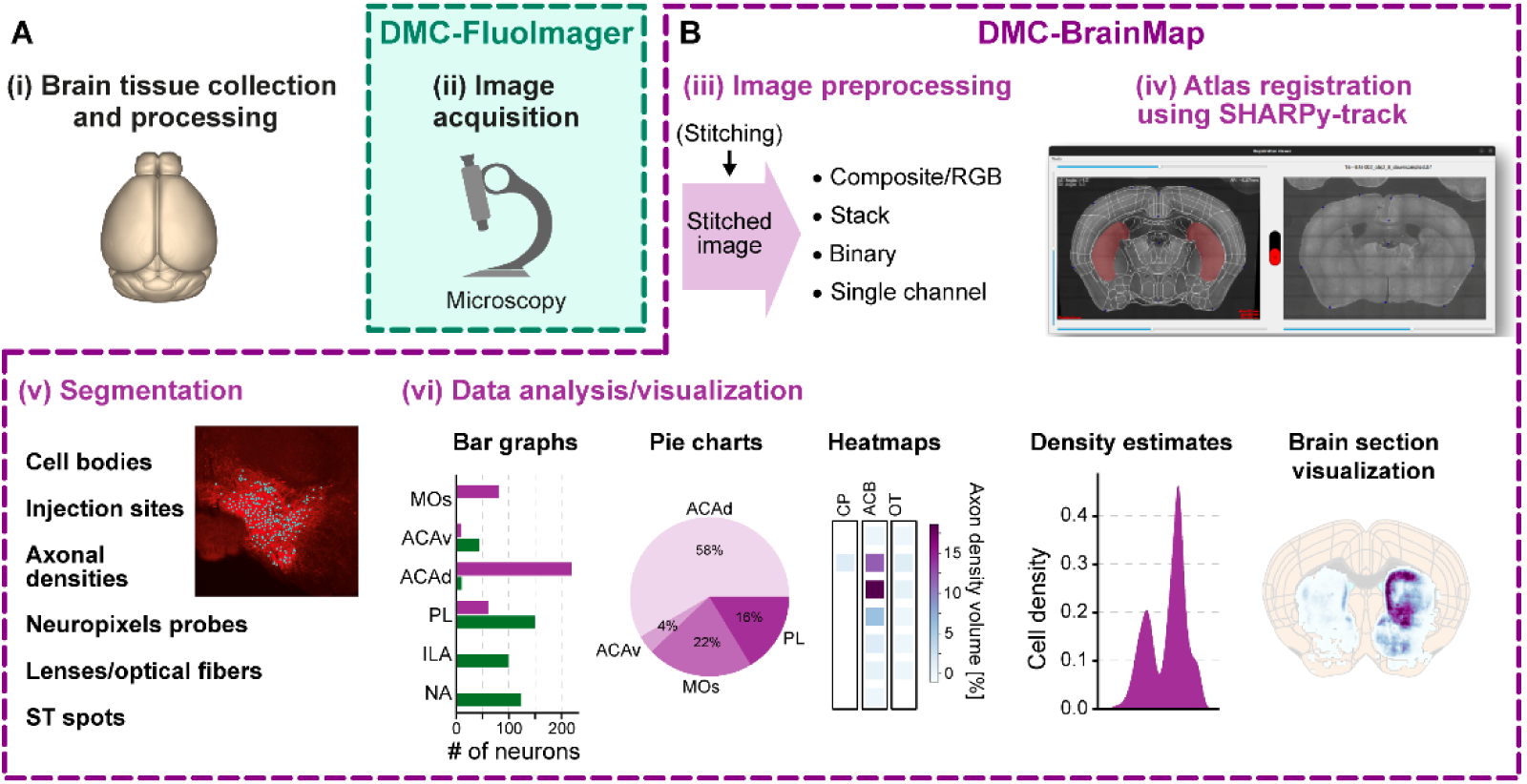
Schematic of the DMC-BrainMap workflow. **(A-B)** The *DMC-BrainMap* workflow provides streamlined processing and whole-brain analysis of anatomical data **(i)**. The pipeline integrates all steps **(iii-vi)** after image acquisition. **(A) (ii)** We provide *DMC-FluoImager* (https://github.com/hejDMC/dmc-fluoimager), a stand-alone solution for automatic imaging of 2D brain sections with a fluorescence microscope. **(B) (iii)** The *DMC-BrainMap* can be used for image preprocessing of stitched images to generate image stacks, composite (RGB), binary, or single channel images. **(iv)** Image registration to a reference atlas is performed using SHARPy-track. **(v)** Various experimental or biological features can be segmented, including cell bodies, injection sites, axonal densities, tracts of Neuropixels probes and optical fibers, as well as ST spots. **(vi)** The anatomical data can be quantified and visualized in many different ways. ACAd, Anterior cingulate area, dorsal part; ACAv, Anterior cingulate area, ventral part; ACB, Nucleus accumbens; CP, Caudoputamen; ILA, Infralimbic area; MOs; Secondary motor cortex; NA, not assessed; OT, Olfactory tubercle; PL, Prelimbic area. Schematic in **(i)** generated using *brainrender* [6].

### Preprocessing of histological images

To start any type of data analysis for an animal for the first time, the user creates a parameter file (*params.json*). This file specifies mandatory information such as the orientation of brain slices (coronal/horizontal/sagittal), acquired fluorophore emission channels, and the reference atlas of choice and optional experimental information (e.g. the specimen’s experimental group/genotype). The *DMC-BrainMap* is compatible with all reference atlases incorporated in the BrainGlobe Atlas API [18] and, thus, enables registration of images from various species and developmental stages. The parameter file keeps a record of analysis steps performed.

The first step in the *DMC-BrainMap* pipeline is image preprocessing. The user can stitch image tiles (a step with integrated padding), or, if working with already stitched images, perform an image padding step (recommended) to align the aspect ratio of acquired images to that of the reference atlas. Various options of batch image processing can be performed on the stitched images (e.g. image contrast adjustment, downsampling) to create single-channel, binarized, or composite (RGB) images, respectively, as well as multichannel image stacks.

### Registration of brain section images to reference atlases using *SHARPy-track*

The *DMC-BrainMap* registers images of histological 2D brain sections into a standardized coordinate system, i.e. the reference atlas selected by the user (Fig 2A). For fast, interactive (immediate feedback) image registration, we developed a tool called *SHARPy-track* (Fig 2B-E). The user starts by specifying the fluorophore emission channel to be used for aligning images of brain sections to planes of the reference atlas. The user thereafter launches a window that will display a plane in the reference atlas (Fig 2B, left) and a brain section image to be registered (Fig 2B, right). The window holds four scrollbars: one for scrolling through and selecting a brain section image, and three for modulating the plane within the reference volume. The user scrolls through the reference volume, following the axis of brain sectioning, to identify the plane matching the brain section image. A common problem when registering histological 2D data is that brains are sectioned at angles slightly deviating from the strictly coronal/sagittal/horizontal reference atlas planes. *SHARPy-track* allows users to correct tissue sectioning angles – the reference plane can be freely rotated (top + left scrollbar) in any direction around the center point of the plane (Fig 2C). For registration of a set of images, the integrated *Registration Helpe*r tool can be used to set the corresponding reference planes for all images at once (Fig 2D). For this, the user assigns an anchor to an arbitrary number of ordered brain section images, e.g. to the first and the last brain section image. The *Registration Helper* tool will interpolate the reference planes of the brain section images between pairs of neighboring anchor images.

**Fig 2.**
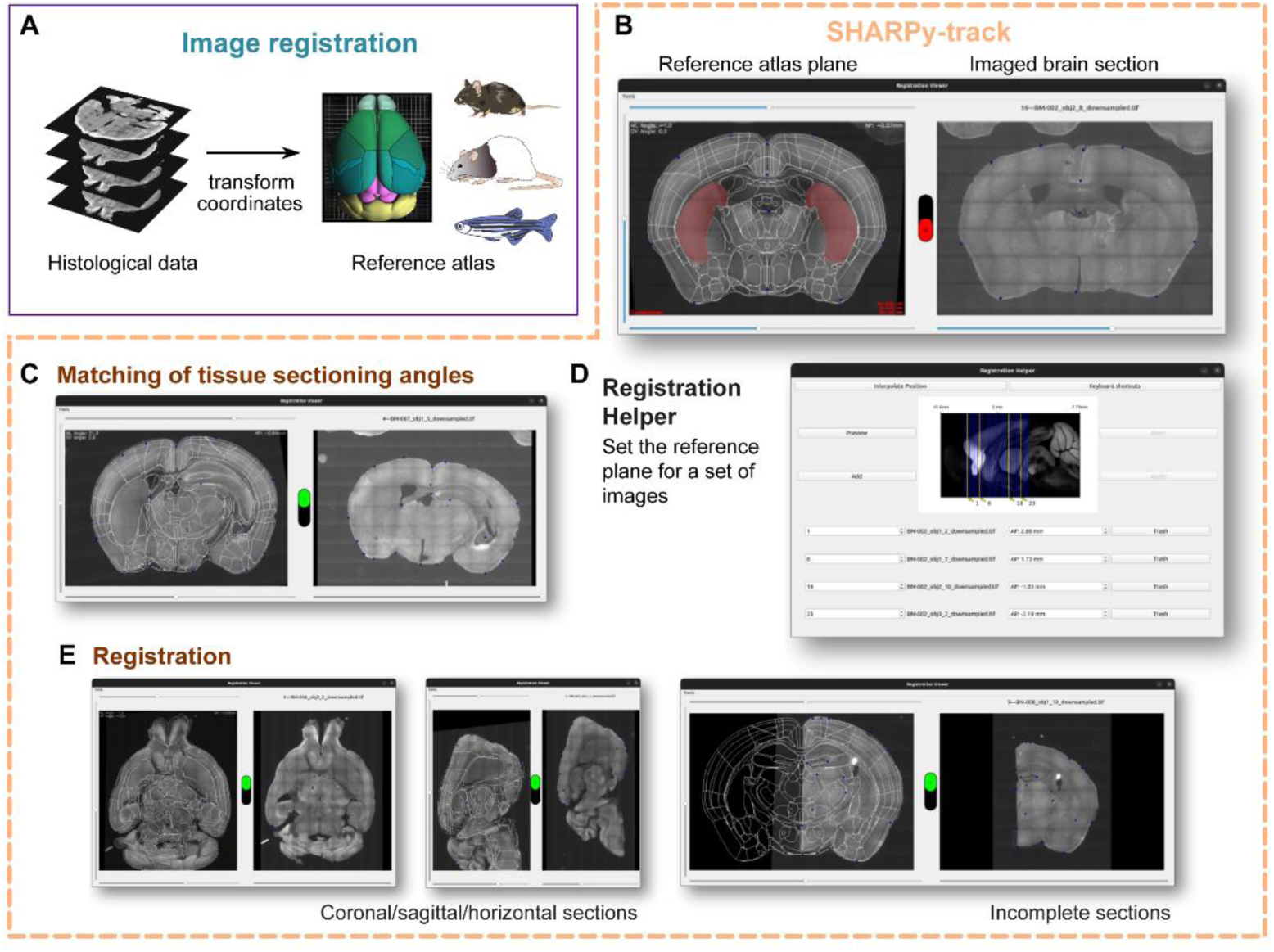
Registration of brain section images to a reference atlas using SHARPy-track. **(A)** The goal of image registration is to anatomically map images of brain sections to a coordinate system (reference atlas). **(B)** The *DMC-BrainMap* pipeline uses the tool *SHARPy-track* for semi-automatic image registration. **(C)** *SHARPy-track* allows free rotation of the reference atlas plane to compensate for tissue sectioning deviating from a strict coronal/sagittal/horizontal plane. **(D)** The *Registration Helper* tool integrated in *SHARPy-track* enables batch setting of reference planes for a set of ordered images. **(E)** *SHARPy-track enables* registration of images of coronal, horizontal, or sagittal brain sections. Images of complete, incomplete, or parts of brain sections can also be registered. Mouse, rat and zebrafish schematic adopted from *scidraw.io*.

To register a brain section image the user manually sets reference points on the reference plane – corresponding (paired) points are automatically generated on the brain section images (the user can easily identify paired points by hovering the cursor over points). The user drags the points on the brain section images to match the location of the reference points. After five (paired) points are set, *SHARPy-track* calculates a perspective transformation matrix to warp the brain section image. The warped image is transparently blended with the reference plane, allowing the user to assess the alignment. An advantage of *SHARPy-track* is that the transformation matrix is calculated in real-time, directly warping the brain section image when the user modifies the position of points. This immediate feedback speeds up the registration process and supports user modifications improving the registration accuracy (Video in Video S1). Importantly, *SHARPy-track* allows users to register images of coronal, sagittal, or horizontal brain sections to a reference atlas, as well as incomplete or parts of brain section images (Fig 2E). To optimize the correspondence between the brain section image and the reference plane, a prediction function keeps track of the latest transformation matrix, allowing SHARPy-track to perform an up-to-date ‘best guess’ in the generation of paired points. Consequently, generating reference points, and adjusting the location of corresponding points, updates the transformation matrix and refines the anatomical precision.

**Video S1. Registration of brain section images to a reference atlas using SHARPy-track.**

### Segmentation and data visualization

Segmentation is the process of assigning labels to distinct features in images. The *DMC-BrainMap* pipeline can segment an array of features, e.g., cell bodies, axonal densities, Neuropixels 1.0 probe tracts, injection sites, and tracts of lenses or optical fibers. In addition, analysis and visualization of ST data is feasible. Importantly, different types of features can be segmented in the same image. The *DMC-BrainMap* pipeline offers a range of options for analysis and visualization of the data (see further below). The data underlying every visualization plot is saved as an individual .csv file, allowing users to run additional, custom analysis or visualization workflows.

### Quantification and visualization of data at the whole-brain level

To demonstrate the utilization of the *DMC-BrainMap* for segmentation and visualization of cell bodies at the whole-brain level, we used rabies virus (RV) tracing in adult wild-type (wt; C57BL/6J) mice to label monosynaptic input to neurons in the dorsomedial and ventromedial prefrontal cortex (dm/vmPFC, respectively; Fig 3A and Fig S1A-B). Segmentation of input neurons, labeled by only RV-eGFP, was performed semi-automatically (Fig S1C). First, the input neurons were automatically segmented using the integrated presegmentation tool (adapted from The Allen Cell and Structure Segmenter ([20]; https://github.com/AllenCell/aics-segmentation workflow). The generated (pre-)segmentation dataset was subsequently manually curated – segmented artifacts were removed and unsegmented neurons were added. Starter neurons (labeled by both helper virus and RV-eGFP; Fig S1C) were manually segmented (see the online user guide for details) and visualized on schematic brain sections, integrating data from several animals (Fig 3B). Quantified starter neurons across the PFC subregions were visualized as pie charts, and in the form of a bar graph (Fig 3C-D). As expected, the locations of starter neurons reflected the injection site in the dmPFC versus the vmPFC, respectively.

**Fig 3.**
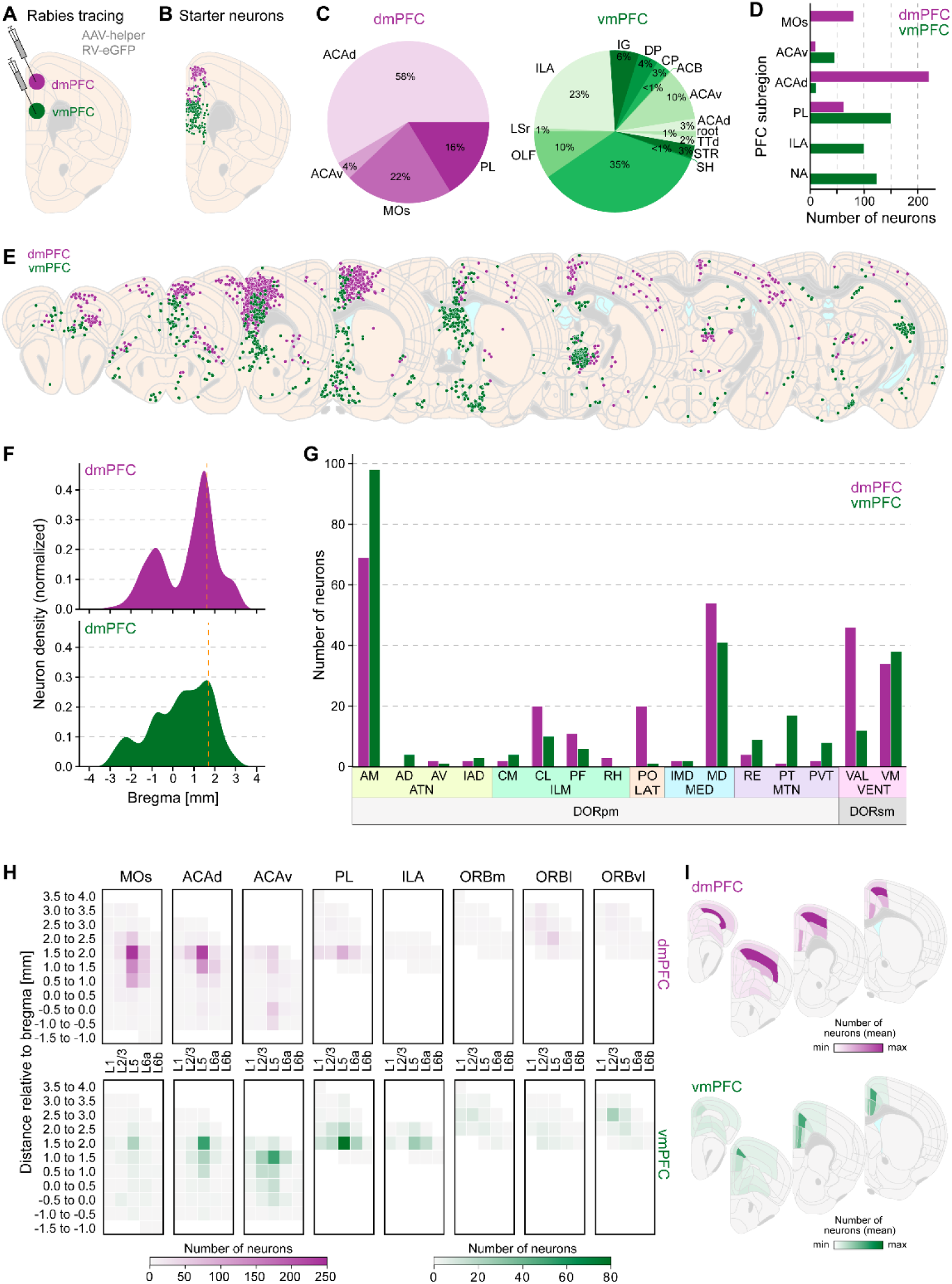
Quantification and visualization of data at the whole-brain level. **(A)** To generate exemplifying data of retrograde, monosynaptic circuit tracing at the whole-brain level, an AAV helper virus cocktail (1:5 AAV5-hSyn-Cre and AAV5-EF1a-DIO-TVA-V5-RG) was injected into the dmPFC (purple) or vmPFC (green) of adult wt mice (n = 1 + 1). Four weeks later RV-eGFP was injected at the same location. **(B)** Segmented starter neurons (V5^+^/RV-eGFP^+^) plotted on a schematic coronal brain section (one hemisphere). Colors as in **(A)**, one dot = one neuron. **(C, D)** Quantification of starter neurons across the subregions of the PFC, visualized as pie charts **(C)** and in a bar graph **(D)**. **(E)** Segmented input neurons (V5^-^/RV-eGFP^+^) plotted on schematic coronal brain sections (anterior to posterior). Monosynaptic input to the dmPFC: purple; to the vmPFC: green. One dot = one neuron. **(F)** Density estimate of input neurons along the AP axis of the brain. **(G)** Bar graph with quantification of the number of input neurons in subregions of the thalamus. **(H)** Heatmaps displaying the laminar distribution of local input neurons along the AP axis of subregions of the PFC. **(I)** Schematic coronal brain sections (anterior to posterior) with heatmaps displaying the distribution of local input neurons across the subregions of the PFC. Included data in **(B, E)**: ± 0.15 mm to the respective brain section. AM, Anteromedial nucleus; AD, Anterodorsal nucleus; ATN, Anterior group of the dorsal thalamus; AV, Anteroventral nucleus of thalamus; CL, Central lateral nucleus of the thalamus; CM, Central medial nucleus of the thalamus; DORpm, Thalamus, polymodal association cortex related; DORsm, Thalamus, sensory-motor cortex related; DP, Dorsal peduncular nucleus; IAD, Interanterodorsal nucleus of the thalamus; IG, Induseum griseum; ILM, Intralaminar nucleus of the dorsal thalamus; IMD, Intermediodorsal nucleus of the thalamus; LAT, Lateral group of the dorsal thalamus; LSr, Lateral septal nucleus, rostral (rostroventral) part; MD, Mediodorsal nucleus of thalamus; MED, Medial group of the dorsal thalamus; MTN, Midline group of the dorsal thalamus; OLF, Olfactory areas; ORBl, Orbital area, lateral part; ORBm, Orbital area, medial part; ORBvl, Orbital area, ventrolateral part; PF, Parafascicular nucleus; PO, Posterior complex of the thalamus; PT, Parataenial nucleus; PVT, Paraventricular nucleus of the thalamus; RE, Nucleus of reuniens; RH, Rhomboid nucleus; SH, Septohippocampal nucleus; STR, Striatum; TTd, Taenia tecta, dorsal part; VAL, Ventral anterior-lateral nucleus of the thalamus; VENT, Ventral group of the dorsal thalamus; VM, Ventral medial nucleus of the thalamus.

**Fig S1.**
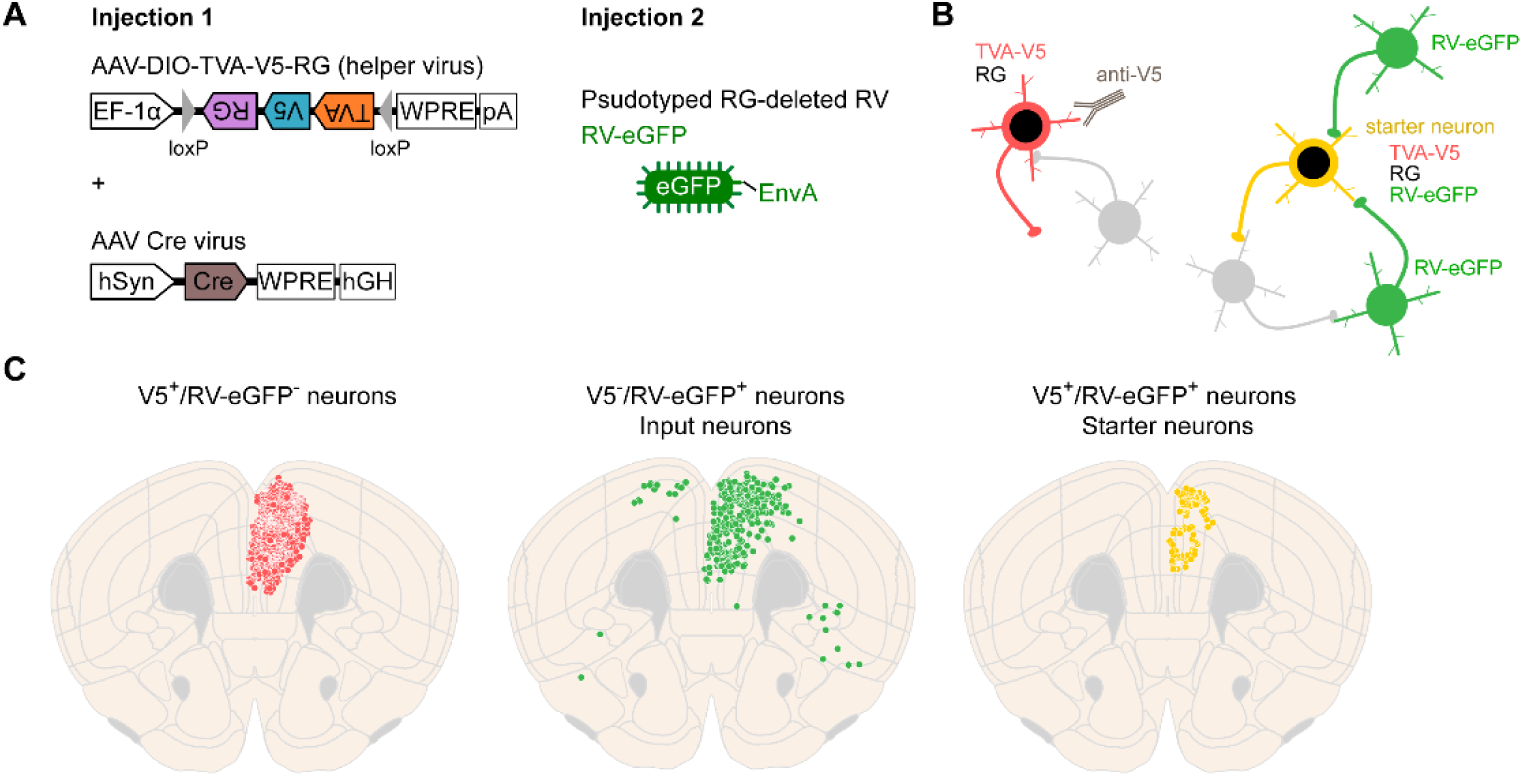
Principle of retrograde rabies tracing of monosynaptic input. **(A)** Injection 1: The helper virus AAV5-DIO-TVA-V5-RG was co-injected with AAV5-hSyn-Cre for pan-neuronal expression of RG and the TVA receptor fused to a V5 tag (TVA-V5), respectively. Injection 2: The RG-deleted RV, pseudotyped with EnvA, expresses eGFP. See [21] for details. **(B)** Starter neurons, giving rise to retrograde transsynaptic labeling of input neurons, are identified by co-expression of TVA-V5 and RV-eGFP (V5^+^/RV-eGFP^+^). Input neurons express only RV-eGFP (V5^-^/RV-eGFP^+^). See [21] for details. **(C)** Schematic coronal brain sections with segmented neurons. Red: V5^+^/RV-eGFP^-^; green: V5^-^/RV-eGFP^+^ = input neurons; yellow: V5^+^/RV-eGFP^+^ = starter neurons. One dot = one neuron. Included data: ± 0.15 mm to the respective brain section.

**Fig S2.**
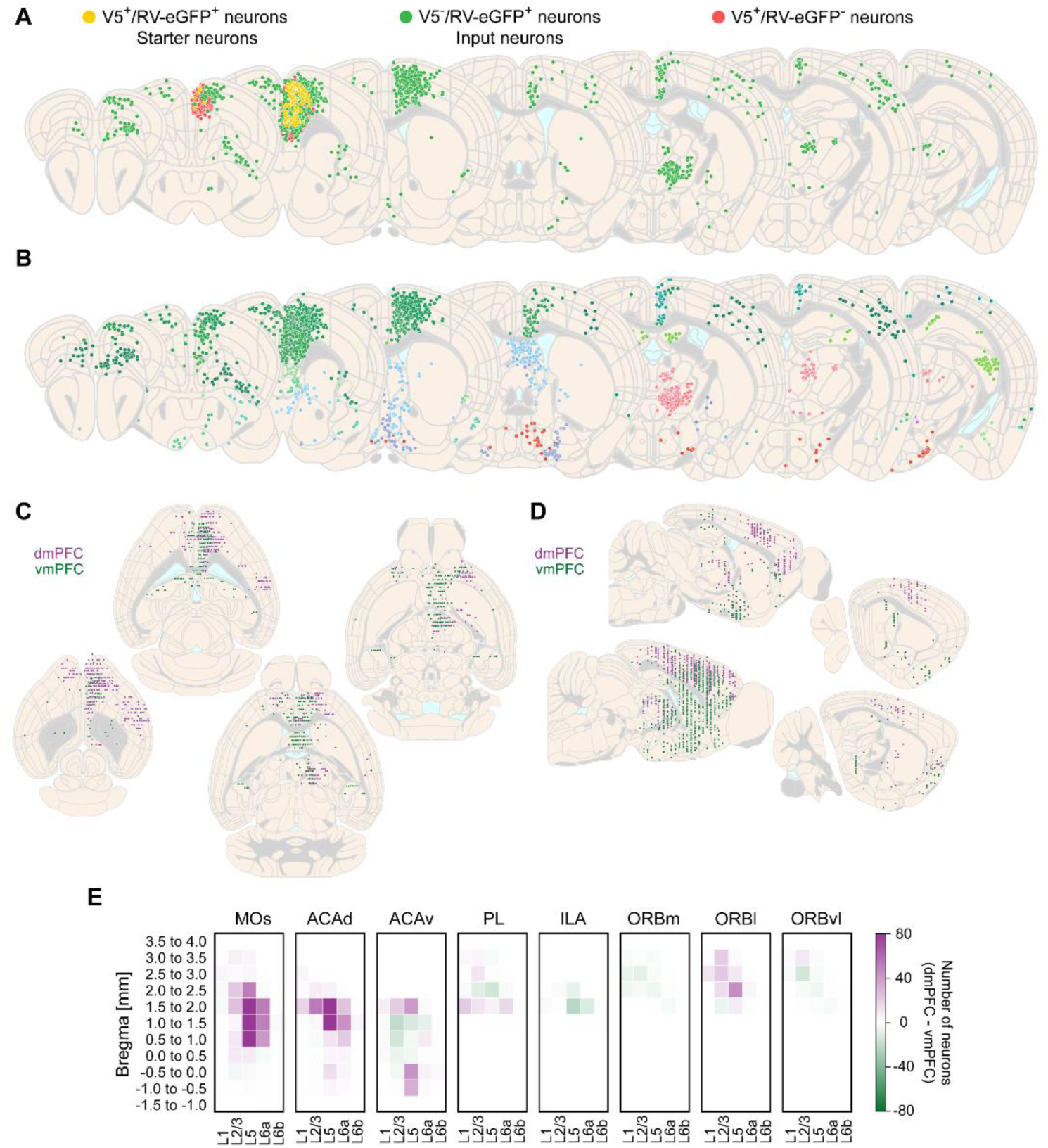
Quantification and visualization of data at the whole-brain level (extended 1/2). **(A)** Segmented neurons plotted on schematic coronal brain sections (anterior to posterior) based on data from one mouse (BM-001; dmPFC-injected). Red: V5^+^/RV-eGFP^-^; green: V5^-^/RV-eGFP^+^ = input neurons; yellow: V5^+^/RV-eGFP^+^ = starter neurons. One dot = one neuron. **(B)** Segmented input neurons (V5^-^/RV-eGFP^+^) plotted on schematic coronal brain sections (anterior to posterior) color-coded according to the Allen Mouse Brain Atlas (CCFv3, [22]). One dot = one neuron. **(C, D)** Segmented input neurons plotted on schematic horizontal (dorsal to ventral) and sagittal (medial to lateral) brain sections. Monosynaptic input to the dmPFC: purple; to the vmPFC: green. One dot = one neuron. **(E)** Heatmap displaying the differential (dmPFC-vmPFC) laminar distribution of local input neurons along the AP axis of the subregions of the PFC. Included data: **(A, B)** ± 0.15 mm, and **(C, D)** ± 0.25 mm to the respective brain section.

**Fig S3.**
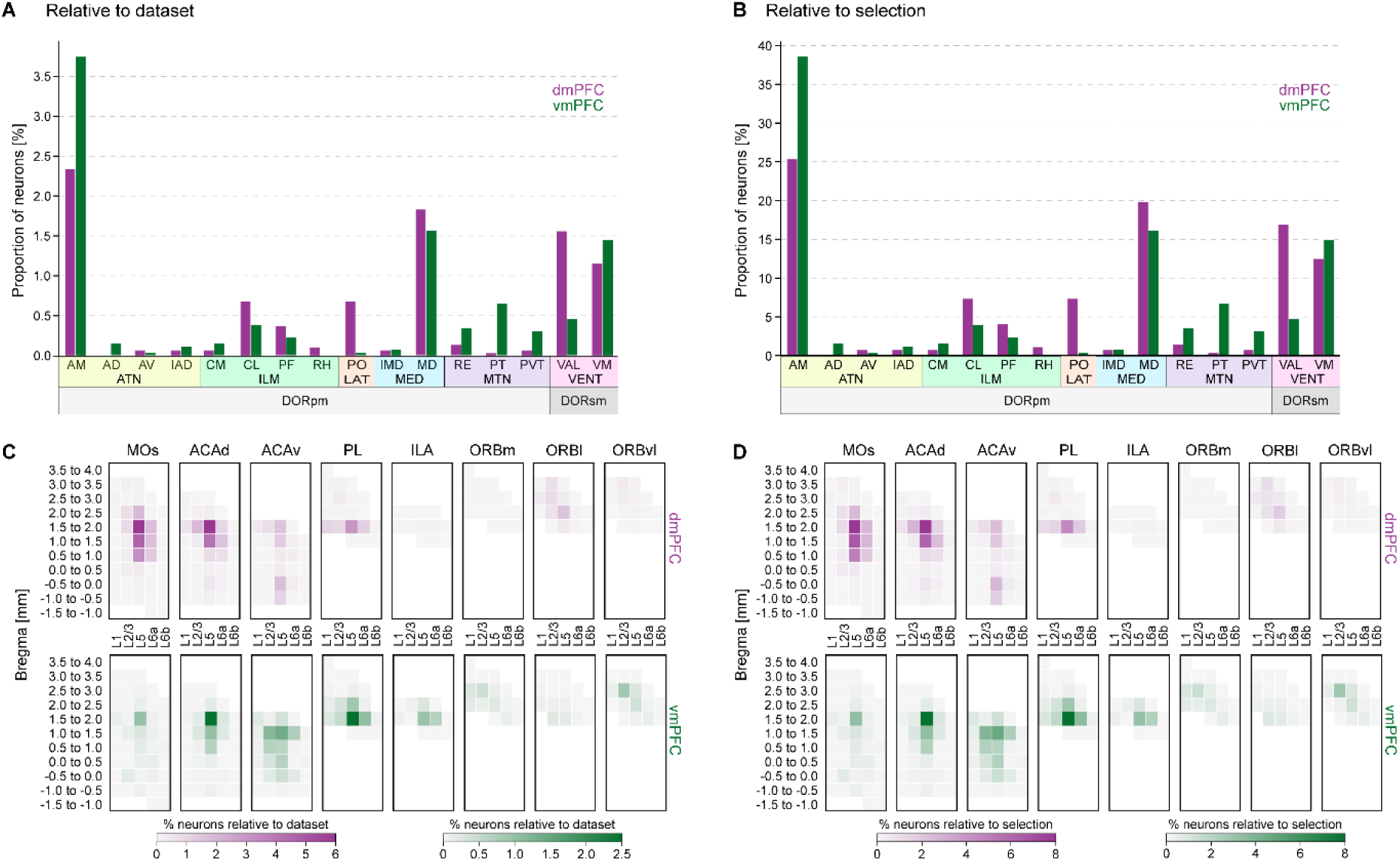
Quantification and visualization of data at the whole-brain level (extended 2/2). (**A, B**) Bar graph with quantification of the relative proportion of input neurons (V5^-^/RV-eGFP^+^) in subregions of the thalamus. Proportions are relative to the total number of all input neurons in the respective dataset (dmPFC (purple) or vmPFC (green; **(A)**), or relative to the total number of input neurons in the selected (i.e., plotted) thalamic subregions in the respective dataset **(B)**. **(C, D)** Heatmaps displaying the relative laminar distribution of local input neurons along the AP axis of the subregions of the PFC. Distributions are relative to the total number of all input neurons in the respective dataset (dmPFC (purple) or vmPFC (green; **(C)**), or relative to the total number of input neurons in the PFC in the respective dataset **(D)**.

Users can compile and visualize segmented features on the whole-brain level on schematic brain sections, classifying the data based on different criteria, e.g., specimen identity, experimental group (Fig 3E), or marker protein expression (Fig S2A). Anatomical features can also be visualized per anatomical annotations in a specific atlas (Fig S2B), including on schematized coronal, horizontal, or sagittal brain sections, irrespective of how the brain slicing was orientated (Fig S2C-D).

Density estimates can be used to exemplify the distribution of neurons along different axes of the brain. *DMC-BrainMap* allows to calculate density estimates along the antero-posterior/dorso-ventral/medio-lateral (AP/ML/DL) axes as well as along a combination of axes. Here, we calculated density estimates of input neurons along the AP axis (Fig 3F). Visualization of the distribution of neurons in selected brain (sub)regions is also straightforward, e.g., in the form of a bar graph (Fig 3G). The absolute number, or proportion of neurons relative to a selected dataset, can be plotted (Fig S3A-B). Heatmaps provide an informative illustration of the distribution of anatomical features in layered structures (Fig 3H and Fig S3C-D), including when plotted on schematic brain sections (Fig 3I).

### *ProbeViewer* for reconstruction and visualization of Neuropixels probe tracts

The *DMC-BrainMap* integrates the tool *ProbeViewer* for reconstructing Neuropixels (1.0) probe tracts, a core functionality of the pipeline (Video in Video S2). Using the 10 μm Allen Mouse Brain Atlas (CCFv3, [22]) or the Waxholm Space (WHS) Atlas of the Sprague Dawley Rat Brain [23] users can anatomically trace the tracts of multiple Neuropixels probes in a single brain, and assign each probe recording site to an anatomical location. For demonstration, we implanted a Neuropixels probe 1.0 labeled with DiO in an adult wt mouse, inserting the probe through the secondary motor cortex (MOs) to the caudoputamen (CP; Fig 4A). The first step is to manually mark the DiO labeled probe tract with reference points in all relevant images. The *objects.Line.best_fit* function of the *scikit-spatial* Python package is thereafter used to reconstruct the probe path based on the set of reference points. The probe insertion depth can be specified by the user or automatically calculated using the reference points. The reconstructed probe tract can be plotted on coronal brain sections using the visualization functions of the *DMC-BrainMap* pipeline (Fig 4B).

**Fig 4.**
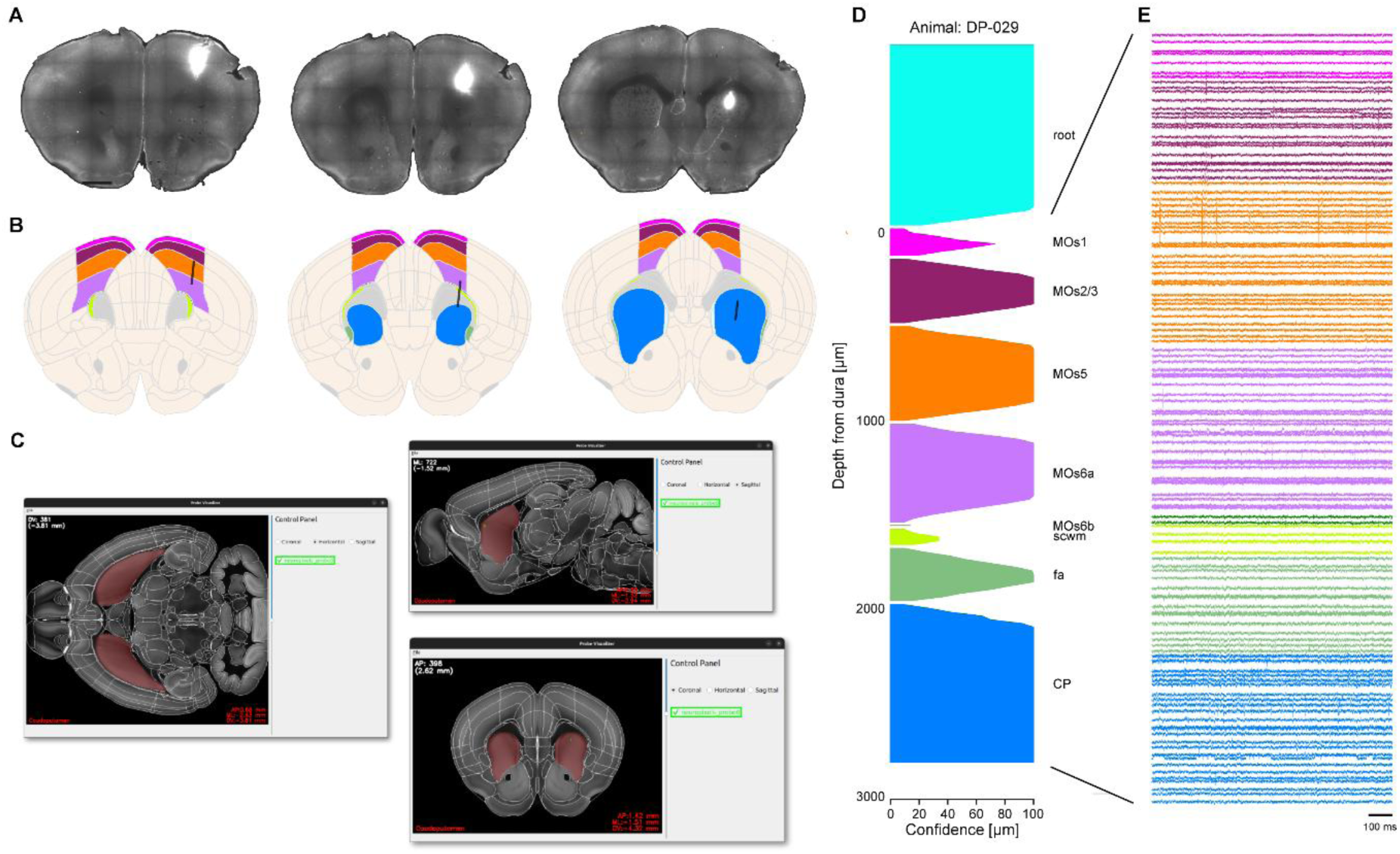
Reconstruction and visualization of Neuropixels probe tracts. **(A)** Coronal brain sections (anterior to posterior) with a Neuropixels probe track labeled with DiO (white). Scale bar: 1 mm. **(B)** Reconstruction of the anatomical localization of the probe in (**A**; black line) based on manual segmentation of the labeled probe tract. Traversed brain regions are custom color-coded using a *DMC-BrainMap* functionality. Included data: ± 0.1 mm to the respective brain section. **(C)** The *ProbeViewer* tool can be used to visually inspect the reconstructed probe tract (green) in different orientations in 3D space. **(D)** Visualization of the brain regions along the reconstructed probe track. The confidence that the brain-region label is correct is indicated by estimation of the distance to the closest brain (sub)region within a 100 μm radius. This confidence is reflected by the thickness along the x-axis. Brain regions color-coded as in **(B). (E)** High-pass filtered electrophysiological traces, color-coded for brain region as in **(B)**. 1 trace = 1 recording channel. fa, corpus callosum, anterior forceps; MOs1, Secondary motor area, layer 1; MOs2/3, Secondary motor area, layer 2/3; MOs5; Secondary motor area, layer 5; MOs6a; Secondary motor area, layer 6a; MOs6b, Secondary motor area, layer 6b; scwm, supra-collosal cerebral white matter.

Using the *ProbeViewer* tool, the user can load the data from multiple probes (from one or more animals) and follow the tracts along the AP, DV, and ML axes of the brain, visually inspecting the reconstructed anatomical paths and their relative positions (Fig 4C). The individual recording sites can be calculated and graphically represented (Fig 4D). The accuracy of the anatomical location of the individual recording sites is indicated by calculation of a confidence metric estimating the distance to the closest next brain (sub)region within a 100 μm radius (see user guide for details). Ultimately, users can integrate the location of the individual recording site in their analysis of neuronal activity signals (Fig 4E).

**Video S2. *ProbeViewer* for visualization of reconstructed Neuropixels tracts**.

### Reconstruction and visualization of injection sites, optical fibers, and axonal densities

The *DMC-BrainMap* can be efficiently used for characterizing the spread of tracers or viral vectors and for validation of optical fiber placement in e.g. optogenetic or imaging experiments. To demonstrate this functionality, we targeted an AAV with pan-neuronal expression of an opsin fused to the fluorophore eYFP to orbital areas (ORB) of adult wt mice (n = 2) and implanted a CM-DiI-labeled optical fiber dorsal to the injection site (Fig 5A). Users can manually segment the anatomical territory holding viral expression using the polygon or polygon lasso objects (see [11,24]) from the *shapes* layer of *napari*. The proportion of segmented pixels across different brain regions can be calculated and visualized as pie charts and bar graphs (Fig 5B-C).

**Fig 5.**
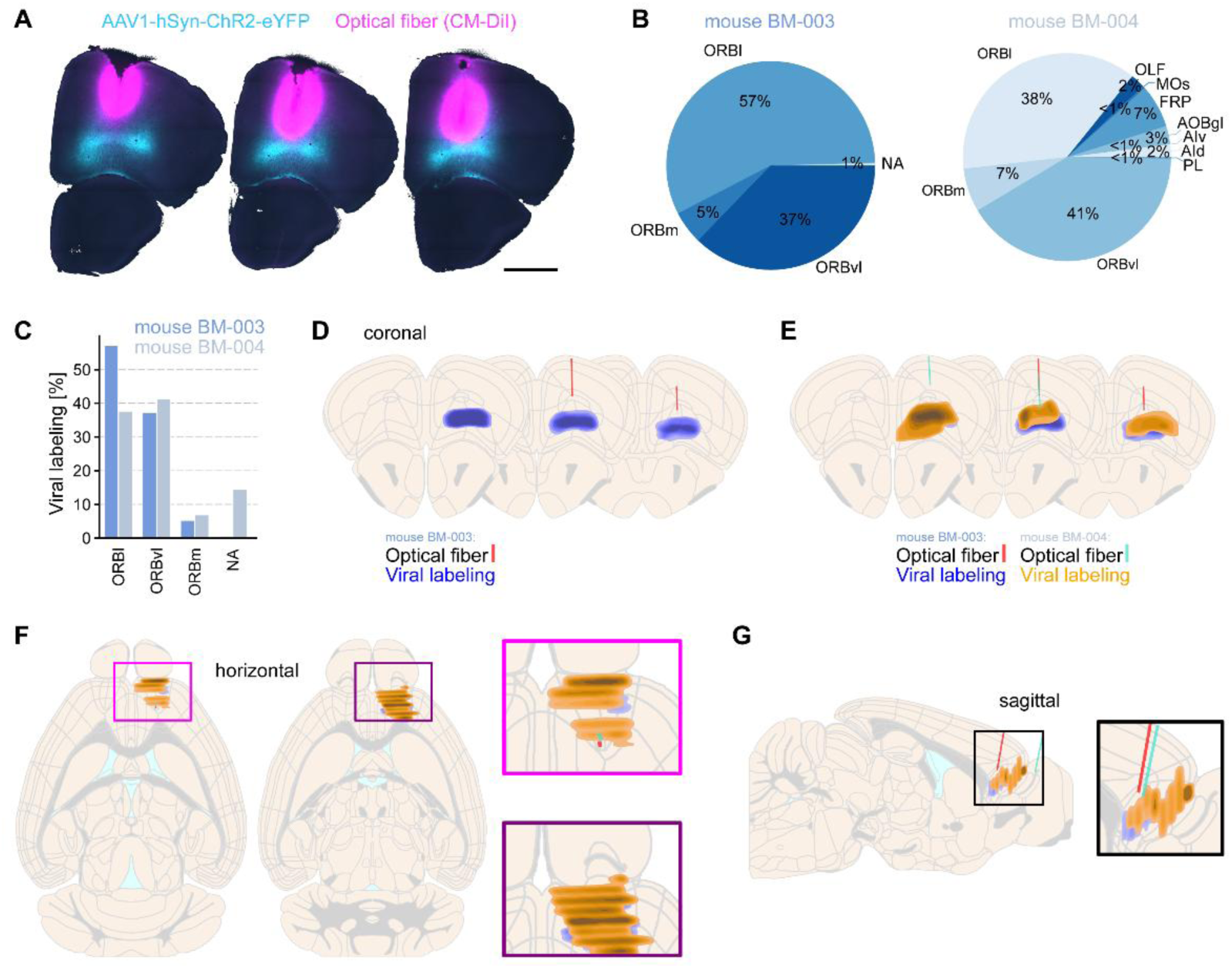
Combined reconstruction of viral labelling and optical fiber tracts. **(A)** To generate exemplifying data for characterizing the spread of viral vectors and validation of optical fiber placement, AAV1-hSyn-ChR2-eYFP (cyan) was targeted to the ORB of adult mice (n = 2). An optical fiber labeled with CM-DiI (pink) was subsequently implanted 0.2 mm dorsal to the injection site. Coronal brain sections (one hemisphere, anterior to posterior) from one of the mice (BM-003). Scale bar: 1 mm. **(B)** Quantification of the viral labeling (manually segmented) across brain regions in the two mice, visualized as pie charts. **(C)** The same data as in **(B)**, visualized as a bar graph with breakdown of the viral labeling in the orbital subregions and all other brain regions (pooled as NA), respectively. **(D)** Schematic coronal brain sections (anterior to posterior) with reconstruction of the anatomical location of the optical fiber (red) and the viral labelling (blue) based on manual segmentation (mouse BM-003). **(E)** Same as **(D)** but with the data for both mice. **(F)** Left: same data as in **(E)**, visualized on schematic horizontal brain sections (dorsal to ventral). Right: magnification of boxes on the left. **(G)** Left: same data as in **(E)**, visualized on a schematic sagittal brain section. Right: magnification of box on the left. Included data: **(D, E)** ± 0.15 mm, **(F)** ± 0.25 mm, and **(G)** ± 0.5 mm to the respective brain section. AId, Agranular insular area, dorsal part; AIv, Agranular insular area, ventral part; AOBgl, Accessory olfactory bulb, glomerular layer; FRP, Frontal pole, cerebral cortex.

The segmentation of the optical fiber tract mirrors the process for Neuropixels probes (see above) and the reconstructed fiber tract can be visualized on schematized brain sections (Fig 5D). Again, different types of segmentation data can be visualized in combination, including data from multiple animals, and the data can be plotted on coronal, horizontal, or sagittal schematized brain sections, irrespective of how the brain slicing was orientated (Fig 5E-G). The output data can be used for experimental evaluation and definition of inclusion/exclusion criteria.

While the anatomical location of neuronal cell bodies is a key parameter in many experiments, assessment of axonal tracts can also be of importance and the *DMC-BrainMap* therefore encompasses analysis of axonal densities. To highlight this functionality, we targeted an AAV with Cre-dependent expression of an opsin fused to the fluorophore mCherry to ventral tegmental area (VTA) in an adult TH-Cre mouse (Fig 6A). The opsin ensures membrane-bound expression of mCherry, including in axons. As a first step, we characterized the viral targeting by segmenting the cell bodies expressing mCherry. As expected, the absolute majority of viral labeling was found in the VTA (Fig 6B). Segmentation of axonal densities using the *DMC-BrainMap* pipeline is done during the preprocessing step, by intensity threshold-based binarization of images, i.e. pixels containing labeled axon segments (high fluorescence: pixel value > threshold) are automatically identified as 1 and pixels without axons (low fluorescence: pixel value < threshold) as 0. This approach is simple and fast, but error-prone. It is also possible to perform the segmentation operation using external tools and software (e.g. [25,26] for volumetric data) and to subsequently integrate the obtained data into the *DMC-BrainMap* workflow. In both cases, the automated segmentation/binarization step is followed by manual curation of the data. The user loads the segmented axonal data and manually removes segmented artifacts. For data analysis and visualization, we used *DMC-BrainMap* to calculate 2D estimates of the axonal densities along both the AP and ML axes of the brain, in our example of axons of TH-positive neurons specifically in the striatum (STR; Fig 6C). Axonal densities in brain (sub)regions can be quantified and plotted e.g., in bar graphs (Fig 6D), and detailed heatmaps can be generated to outline axonal densities along the AP axis with brain region specificity (Fig 6E). Alternatively, axonal densities can be visualized on schematized brain sections (coronal, horizontal, or sagittal orientation; Fig 6F-H).

**Fig 6.**
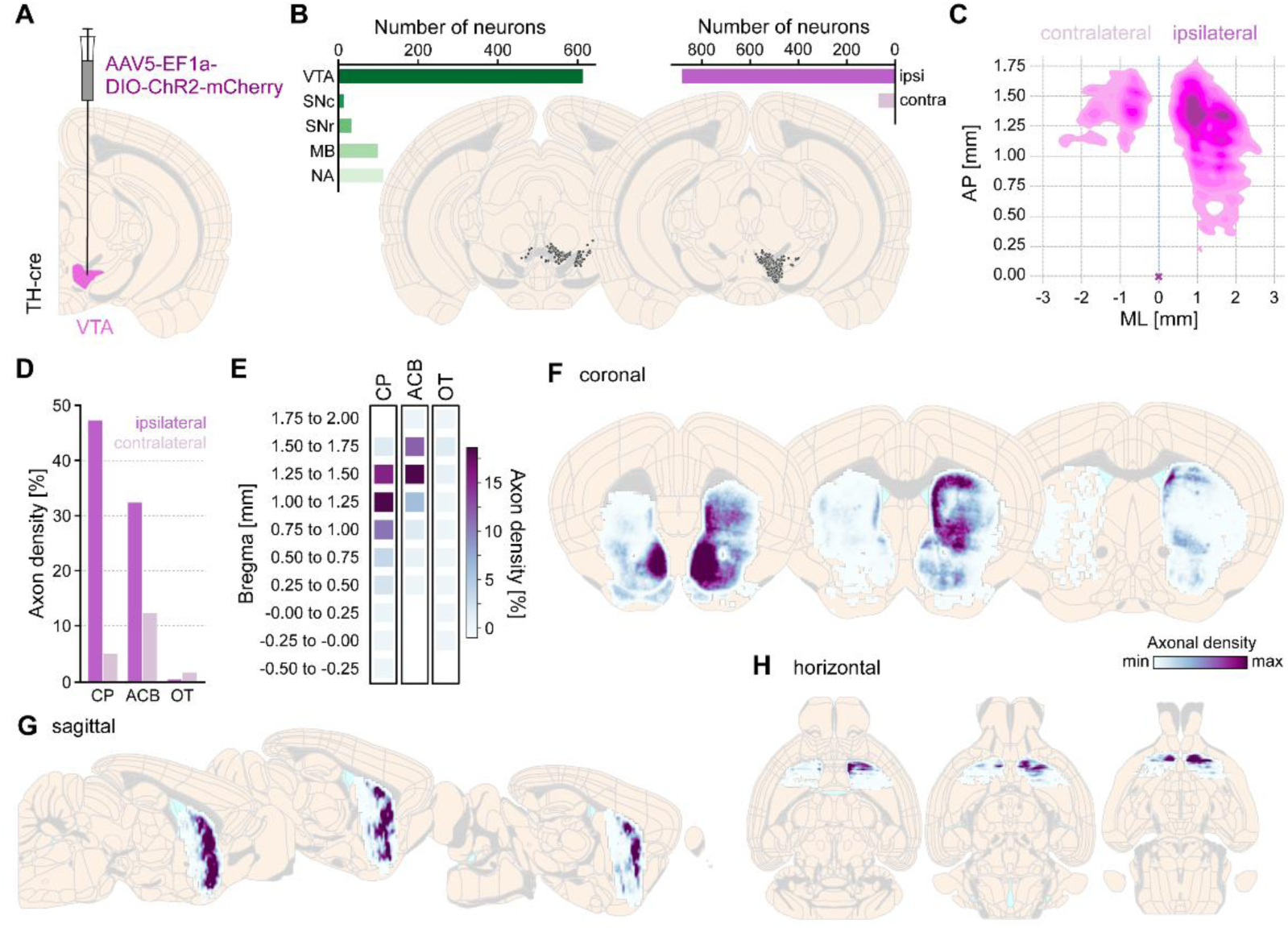
Quantification of axonal densities. **(A)** To generate exemplifying data for assessment of axonal densities, AAV-EF1a-DIO-ChR2-mCherry was unilaterally targeted to the VTA (pink) in an adult TH-Cre mouse (n = 1). **(B)** Segmented mCherry-positive neurons plotted on schematic coronal brain sections. One dot = one neuron. Inlets: Quantification of labeled neurons across brain regions (left) and across the two hemispheres (relative to the injection site; right), plotted as bar graphs. **(C)** Axonal densities were segmented by threshold-based binarization of images and manually curated. 2D plot of the axonal density (Gaussian kernel) along the AP and ML axes of the STR. Purple cross: bregma; dashed blue line: midline. **(D)** Bar graph with quantification of axonal density in subregions of the STR, ipsi-versus contralateral to the injection. **(E)** Heatmaps displaying the axonal density along the AP axis in subregions of the STR. **(F, G, H)** Axonal densities in the STR, plotted on schematic coronal (**F**; anterior to posterior), sagittal (**G**; medial to lateral), and horizontal (**H**; dorsal to ventral) brain sections. Included data: **(B)** ± 0.1 mm, **(F)** ± 0.15 mm, **(G)** ± 0.5 mm, and **(H)** ± 0.25 mm to the respective brain section.

### Analysis and visualization of whole-brain ST data

Sequencing directly in sectioned tissue provides data on the spatial organization of gene expression [27,28]. *DMC-BrainMap* accommodates analysis and visualization of ST data, and we here used a publicly available dataset (https://www.molecularatlas.org/; [29]) for proof-of-concept demonstration. The dataset holds adult wt mouse whole-brain ST data (75 coronal sections, one hemisphere). As a first step, we registered the dataset’s Hematoxylin-Eosin (HE) stained sections to the 10 μm Allen Mouse Brain Atlas (CCFv3, [22]) to map the anatomical location of each spot of the ST array (Fig 7A-B). The expression of single genes can be visualized on schematized brain sections in three different ways, here we used the (normalized) expression of the Slc17a7 gene encoding the vesicular glutamate transporter (Vglut1) to exemplify this; the expression level in individual ST spots can be color-coded, or brain (sub)regions can be color-coded based on the average, normalized gene expression (Fig 7C-D). As an alternative, the pipeline can generate a Voronoi tessellation of the reference atlas to visualize expression levels on brain sections (Fig S4A).

**Fig 7.**
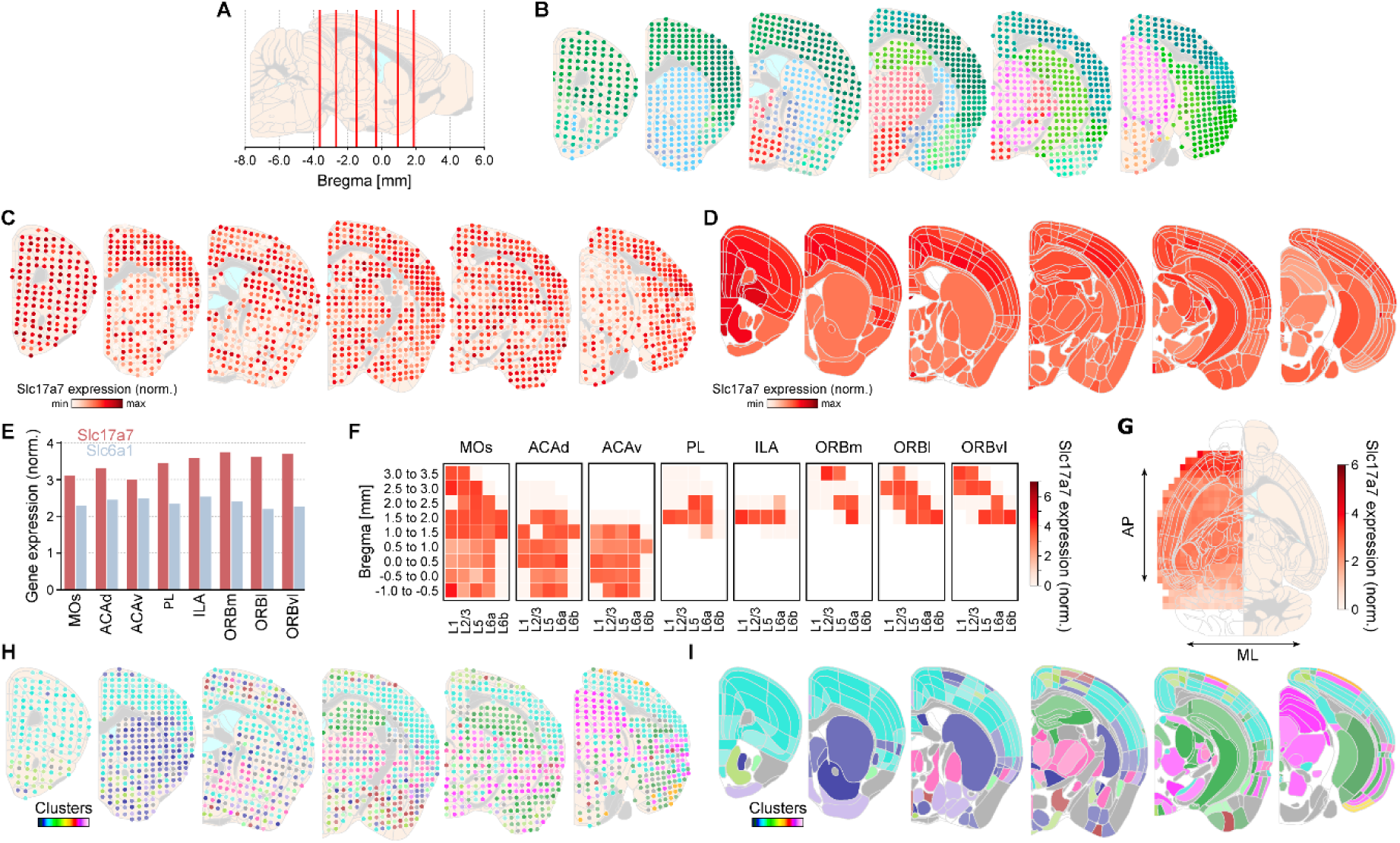
Analysis and visualization of whole-brain ST data. For demonstrating *DMC-BrainMap’s* functionality to analyze whole-brain ST data, a publicly available adult mouse dataset (75 coronal sections, one hemisphere, https://www.molecularatlas.org/; [29]) was registered to the Allen Mouse Brain Atlas (CCFv3, [22]). **(A)** Schematic sagittal brain section with the AP location of six representative brain sections (6/75; red vertical lines) displayed in **(B-D, H-I)**. **(B)** Schematic coronal brain sections (one hemisphere; anterior to posterior) with spots color-coded according to the Allen Mouse Brain Atlas. One dot = one ST spot. **(C)** As **(B)**, but spots color-coded according to Slc17a7 expression (normalized). **(D)** Schematic coronal brain sections (one hemisphere; anterior to posterior) with brain regions color-coded according to Slc17a7 expression (normalized). **(E)** Bar graph with quantification of Slc17a7 and Slc6a1 expression (normalized), respectively, in subregions of the PFC. **(F)** Heatmap displaying the laminar Slc17a7 expression (normalized) along the AP axis of the subregions of the PFC. **(G)** Left: Horizontal brain section (one hemisphere) with heatmap displaying the Slc17a7 expression (normalized) along the AP and ML axes. Right: schematic horizontal brain section (one hemisphere). **(H)** As **(B)**, but spots color-coded according to cluster identity. **(I)** As **(D)**, but brain regions color-coded according to the dominant cluster (‘winner-takes-all’). Included data: **(B, C)** ± 0.0 mm, and **(D, I)** ± 0.05 mm to the respective brain section.

**Fig S4.**
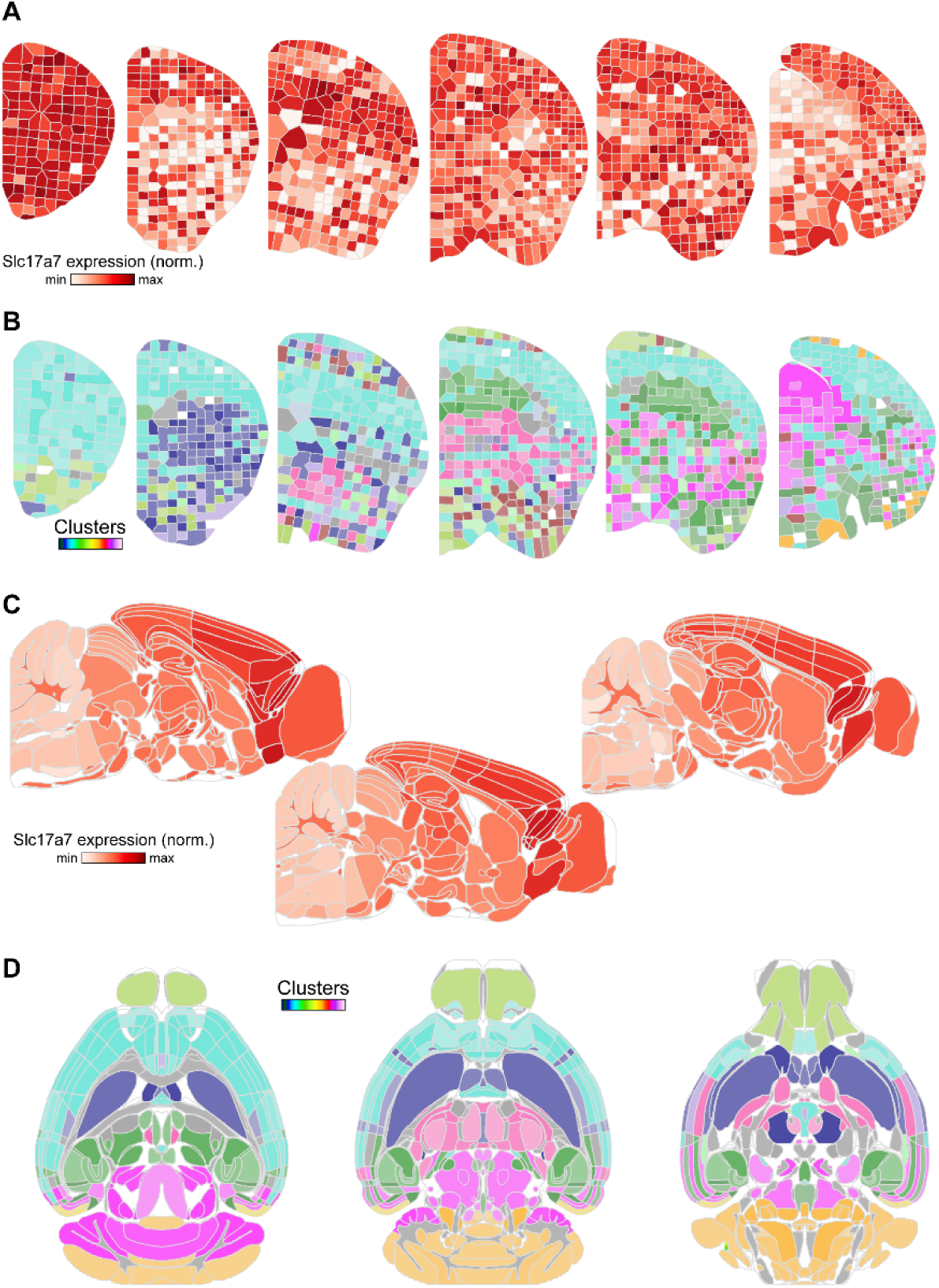
Analysis and visualization of whole-brain ST data (extended). **(A)** Schematic coronal sections (one hemisphere, anterior to posterior) displaying a Voronoi tessellation of Slc17a7 expression (normalized). **(B)** As **(A)**, but with Voronoi tessellation of cluster identity. **(C)** Schematic sagittal brain sections (medial to lateral) with brain regions color-coded according to Slc17a7 expression (normalized). **(D)** Schematic horizontal brain sections (dorsal to ventral) with brain regions color-coded according to the dominant cluster (‘winner-takes-all’). Included data: (A, B) ± 0.05 mm, and (C, D) ± 0.25 mm to the respective brain section.

Quantification of the (normalized) expression levels of a set of genes in selected brain regions is possible by using the bar graph functionality of the pipeline (Fig 7E). Likewise, users can use the heatmap functionality to visualize the (normalized) expression of a gene based on a combination of anatomical parameters, e.g., brain (sub)regions, layers, and an axis of the brain (Fig 7F). Density estimates of a gene’s expression along two axes of the reference atlas, plotted on schematized brain sections, provide an informative illustration of the data at the whole-brain level (Fig 7G).

Clustering of gene expression levels in ST spots is a common strategy to identify molecularly distinct subregions in the sequenced tissue [29]. This type of data can also be visualized using the *DMC-BrainMap* pipeline. The ST spots can be color-coded by their cluster identity, or brain (sub)regions color-coded by the majority of clusters present (‘winner-takes-all’; Fig 7H-I). As for gene expression, a cluster identity-based Voronoi tessellation of the reference atlas is also possible (Fig S4B). As for any other anatomical feature, gene expression or clustering data can be visualized on coronal, horizontal, or sagittal schematized brain sections, irrespective of how the brain slicing was orientated (Fig S4C-D).

## Discussion

### Streamlining quantitative, whole-brain anatomical data analysis

We here introduce the *DMC-BrainMap*, a *napari* plugin for quantitative image analysis and visualization of whole-brain anatomical data across species. From the plethora of anatomical data analysis tools released in recent years (Table in S1 Table), three features set the *DMC-BrainMap pipeline* apart. First, while several tools provide excellent solutions to individual steps in anatomical data analysis [10–16,30], execution of the whole workflow generally demands integration of several tools. The *DMC-BrainMap* provides users with a ’one software solution’ covering all steps from image preprocessing to data visualization. Second, tools like the WholeBrain package [13] and HERBS [11] cover most steps in anatomical data analysis but are limited to analysis of tissue from mice (WholeBrain, HERBS) and rats (HERBS; both species) – the *DMC-BrainMap* pipeline, through its use of the BrainGlobe Atlas API [18], allows users to analyze data obtained across a range of model animals including mice, rats and zebrafishes at developmental and adult stages as well as Mexican cavefishes, lemurs, Axolotls and prairie voles. In addition, a range of mouse brain atlases are included in the BrainGlobe Atlas API. *DMC-BrainMap* provides increased flexibility and versatility also regarding the anatomical features to be segmented (cell bodies, probe/optical fiber tracts, injection sites, axonal densities, gene expression, etc) which is not possible in existing solutions without extensive customization and programming [10]. Thirdly, the *DMC-BrainMap* pipeline is intuitive and easy to use, and installed and run via a GUI, eliminating the requirement of programming skills. The *DMC-BrainMap* is completely written in Python, eliminating the need for software licenses while achieving cross-platform compatibility – the pipeline works for Windows, Mac, and Linux operating systems. All analyzed data is exported in a standardized and organized format, enabling users with programming skills to conduct further sophisticated analysis. Furthermore, data generated by *DMC-BrainMap* is readily formatted to be imported into *brainrender* workflows [6], enabling users to generate e.g. visually appealing 3D renderings of their data.

### Limitations and future improvements of *DMC-BrainMap*

We intentionally developed the *DMC-BrainMap* to allow for manual curation of image registration and anatomical feature segmentation, giving users versatile control over the performed operations. However, manual operations are time-consuming compared to automated approaches. In recent years, approaches such as DeepSlice [31] have significantly improved automatic image registration, with the potential to dramatically reduce the user’s time investment. Intended developments of the *DMC-BrainMap* pipeline include automated options for image registration and anatomical feature segmentation (e.g. axonal densities). In line with this, the *DMC-BrainMap* is fully open-source, enabling the pipeline to be a community-based tool open to iterative improvement and expansion by the scientific community.

## Methods

### Animals

All procedures and experiments on animals were performed according to the guidelines of the Stockholm Municipal Committee for animal experiments and the Karolinska Institutet in Sweden (approval number 7362-2019). See Table in Table 1 for a complete list of animals. Animals were housed in groups with up to four animals per cage, in a temperature (23 °C) and humidity (55%) controlled environment in standard cages on a 12:12 h light/dark cycle with *ad libitum* access to food and water.

**Table 1.** List of animals.

| Animal ID | Genotype | Implant/virus | Coordinates | Data shown |
| --- | --- | --- | --- | --- |
| BM-006 | wt<br>(C57BL/6J) | - | - | Fig 2A, E |
| BM-007 | wt<br>(C57BL/6J) | - | - | Fig 2D |
| BM-005 | wt<br>(C57BL/6J) | - | - | Fig 2E |
| BM-008 | wt<br>(C57BL/6J) | - | - | Fig 2F |
| BM-001 | wt<br>(C57BL/6J) | pENN.AAV5.hSyn.Cre.WPRE.hGH<br>(addgene: 105553; titre:<br>2.1x10<sup>13</sup>; 50 nl)<br>+ AAV5-EF1a-DIO-TVA-V5-RG-WRPE-hGHpA - (virovek #17-042,<br>titer:1.2x10<sup>13</sup>; 250 nl)<br>> 4 weeks:<br>EnvA-dG-eGFP-RV ([21]; 800 nl) | AP: 1.7 mm<br>ML: 0.4 mm<br>DV: 1.0 mm | Fig 3B-H<br>Fig S2D<br>Fig S3 (*all panels*) |
| BM-002 | wt<br>(C57BL/6J) | pENN.AAV5.hSyn.Cre.WPRE.hGH<br>(addgene: 105553; titre:<br>2.1x10<sup>13</sup>; 50 nl)<br>+ AAV5-EF1a-DIO-TVA-V5-RG-WRPE-hGHpA - (virovek #17-042,<br>titer:1.2x10<sup>13</sup>; 250 nl)<br>> 4 weeks: | AP: 1.7 mm<br>ML: 0.4 mm<br>DV: 1.6 mm | Fig 1<br>Fig 2B, C<br>Fig 3B-H<br>Fig S3C-H |
|  |  | EnvA-dG-eGFP-RV ([21]; 800 nl) |  |  |
| DP-029 | wt<br>(C57BL/6J) | Neuropixels probe 1.0 | AP: 1.8 mm<br>ML: 1.5 mm<br>DV: 2.5 mm | Fig 4 (*all panels*) |
| BM-003 | wt<br>(C57BL/6J) | Optical fiber +<br>AAV1-hSyn-ChR2-eYFP (addgene:<br>26973; titre: 2.4x10<sup>13</sup>; 200 nl) | AP: 2.6 mm<br>ML: 1.2 mm<br>DV: 1.4/6<br>mm | Fig 5 (*all panels*) |
| BM-004 | wt<br>(C57BL/6J) | Optical fiber +<br>AAV1-hSyn-ChR2-eYFP (addgene:<br>26973; titre: 2.4x10<sup>13</sup>; 200 nl) | AP: 2.6 mm<br>ML: 1.2 mm<br>DV: 1.4/6<br>mm | Fig 5B-G |
| DP-117 | TH-Cre<br>(Tg(TH-Cre)12Gsat) | AAV5-EF1a-DIO-ChR2-mCherry<br>(200 nl) | AP: -3.0 mm<br>ML: 0.5 mm<br>DV: 4.2 mm | Fig 1<br>Fig 6 (*all panels*) |

AP, antero-posterior; DV, dorso-ventral; ML, medio-lateral; wt, wild-type

## Surgical procedures

### General surgical procedures and post-operative care

The animals were deeply anesthetized with isoflurane in oxygen (4 % for induction, 1-2 % for maintenance), fixed in a stereotaxic frame (Harvard Apparatus), and buprenorphine (0.1 mg/kg s.c.) was thereafter injected. Lidocaine (4 mg/kg s.c.) was injected locally before skin incision. An ocular ointment (Bepanthen) was applied over the eyes, and the body temperature was maintained at 37 °C using a heating pad. After surgery (details below), animals were injected with carprofen (5 mg/kg s.c.) and returned to their homecage. An additional dose of carprofen was delivered 18-24 h after surgery.

### Intracerebral virus injection

For virus injections, an incision into the skin overlying the skull was made and the skin was carefully moved aside. A small craniotomy (0.5 mm diameter) was drilled over the target coordinates (AP relative to bregma, DV relative to dura, ML relative to midline; Table in Table 1). The virus was delivered by a glass capillary attached to a motorized Quintessential Stereotaxic Injector (Stoelting) at a rate of 30-50 nl/min. The capillary was held in place for 5 min before and 10 min after the injection. The incision was closed with stitches (Ethicon). For retrograde tracing, pseudotyped rabies virus (RV) was injected 4 weeks after the first surgery at the same coordinates as the helper AAV injection. General surgical procedures and post-operative care were followed.

### Optical fiber implant surgery

For optical fiber implant surgeries, the skin overlying the skull was removed and the bone was gently cleaned. Custom-build optical fibers (Thorlabs) were labeled with CM-DiI (V22888, Thermo Fisher) and implanted immediately after intracerebral virus injection. Optical fibers were slowly lowered into the existing craniotomy and secured 0.2 mm dorsal to the viral injection site using UV-curable dental cement (Pluline). General surgical procedures and post-operative care were followed.

### Head-post implant surgery and habituation

For head-post implant surgeries, the skin overlying the skull was removed and the bone was gently cleaned. A thin layer of super glue (Loctite) was applied to the skull and a lightweight metal head-post was fixed with super glue and UV-curable dental cement (Pluline). For Neuropixels recordings, a chamber (4 mm diameter) was made using UV-curable dental cement centered around the coordinates for probe insertion. General surgical procedures and post-operative care were followed. Following 7 days of surgery recovery, the mouse was handled and progressively habituated to the head-fixation procedure over 3 days by increasing the head-fixation time from 15 min to 1 h.

### Neuropixels recordings

For acute recordings, two small craniotomies (< 0.5 mm diameter) were opened >3 h before the experiments. One craniotomy was made above the targeted probe insertion site, and one craniotomy was made postero-lateral of bregma and used for reference electrode placement.

General surgical procedures and post-operative care were followed. The open craniotomies were covered with Silicone sealant (Kwik-Cast, WPI), and the mouse was returned to its home cage for recovery. For the recordings, the mouse was head-fixed and the DiO-labeled (V22886, Thermo Fisher) Neuropixels probe was gradually lowered (∼20 μ s−1) into the brain until reaching the target coordinates using a micromanipulator (uMp-4, Sensapex). The electrode reference was connected to a silver wire positioned over the pia in the second craniotomy, using a separate micromanipulator. The Neuropixels probe was allowed to sit in the brain for 20–30 min before the recordings started. The spike band data were digitized with a sampling frequency of 30 kHz with gain 500, transferred to the data acquisition system (a PXIe acquisition module PXI-Express chassis: PXIe-1071 and MXI-Express interface: PCIe-8381 and PXIe-8381; National Instruments) and written to disk using SpikeGLX (Bill Karsh, Janelia). The spike band data were filtered between 0.3 and 10 kHz and amplified (see [32,33] for details).

## Tissue collection and processing

### General procedure

For perfusions, the animals were deeply anesthetized with pentobarbital and transcardially perfused with 0.1 M phosphate buffered saline (PBS) followed by 4% formaldehyde (FA, VWR) in 0.1 M PBS. The perfused brain was removed from the skull and postfixed in 4% FA in 0.1 M PBS at 4 °C for 16 h. The brains were thoroughly washed in 0.1 M PBS and thereafter sectioned (50-100 μm thickness) using a vibratome (Leica VT1000, Leica Microsystems) and mounted on glass slides (Superfrost Plus, Thermo Scientific).

### Immunofluorescence staining

The mounted brain sections were permeabilized with TBST (0.3 % TritonX-100 in Tris-buffered saline (42 mM Trizma hydrochloride (Sigma-Aldrich), 8 mM Trizma base (Sigma-Aldrich), 120 mM NaCl (Merck) in demineralized H2O; TBS) for 1 h, blocked with 10% normal donkey serum in TBST for 1 h, and thereafter incubated with primary antibodies (1:500, Ch-α-V5, abcam (ab9113)) in TBST at room temperature for 12–24 h. The sections were thereafter washed three times in TBST and incubated with a species-specific fluorophore-conjugated secondary antibody (1:500, Cy3-α-Ch, Jackson (703-165-155)) in TBST for 4 h. The sections were thereafter consecutively washed with TBST, TBS, and 0.1 M PBS (10 min each) and stained with 4′,6-Diamidine-2′-phenylindole dihydrochloride (DAPI; Sigma-Aldrich, #10236276001). All sections were coverslipped (Thermo Scientific) using Mowiol 4-88 (Sigma-Aldrich, #81381).

## Data acquisition and analysis

### Fluorescent microscopy

Tiled images were semi-automatically acquired at magnification ×10 (0.64 µm/pixel) using a Leica DM6000B fluorescent microscope with a Hamamatsu Orca-FLASH 4.0 C11440 digital camera at 16-bit depth resolution using the *DMC-FluoImager* script (https://github.com/hejDMC/dmc-fluoimager).

### Mapping of anatomical and histological features to reference atlases

If not stated otherwise, *DMC-BrainMap* was used for all data analysis and visualization. Figures were assembled in Inkscape and Adobe Illustrator. The *DMC-BrainMap pipeline* is entirely written in Python (3.10) using the following packages: aicsimageio (4.14.0), aicspylibczi (3.1.2), aicssegmentation (0.5.3), bg_atlasapi (1.0.2), distinctipy (1.3.4), imagecodecs (2024.1.1), magicgui (0.8.1), matplotlib (3.8.3), mergedeep (1.3.4), natsort (8.4.0), numpy (1.26.4), opencv-python (4.9.0.80), pandas (2.0.1), qtpy (2.4.1), scikit-image (0.22.0), scikit-learn (1.4.1.post1), scikit-spatial (7.2.0), seaborn (0.12.2), shapely (2.0.1), tifffile (2023.2.28). The software was tested on Windows, Mac (both Intel and Apple Silicon), and Linux operating systems. The software is open-source, and available online (https://github.com/hejDMC/napari-dmc-brainmap). Detailed installation guides and documentation are provided online. All data was deposited on *figshare* (https://doi.org/10.6084/m9.figshare.28429478).

## Notes

https://doi.org/10.6084/m9.figshare.28429478

### Competing Interest Statement

The authors have declared no competing interest.

