## Supplementary material for "DMC-BrainMap - an open-source, end-to-end tool for multi-feature brain mapping across species": Table S1

| Tool | Year | Software | OS | Species | Tissue | Registration | Segmentation | Visualization | Additional notes | Reference | Source code |
|---|---|---|---|---|---|---|---|---|---|---|---|
| aMAP | 2016 | Python | Linux, MacOS | Mouse | 3D | auto | yes | no | discontinued, precursor to brainreg | DOI | https://github.com/SainsburyWellcomeCentre/amap_python |
| ClearMap | 2016 | Python | Linux, MacOS | Mouse | 3D | auto | auto (cells and blood vessels) | yes | - | DOI | https://github.com/ChristophKirst/ClearMap2 |
| brain-mapping | 2018 | Matlab | Linux | Mouse | 2D | auto | no | yes | - | DOI | https://sites.google.com/view/brain-mapping |
| WholeBrain | 2018 | R | Linux, MacOS, Windows | Mouse | 2D/3D | auto with curation | auto (cells/axons) | yes | - | DOI | https://github.com/tractatus/wholebrain |
| SHARP-Track | 2018 | Matlab | Linux, MacOS, Windows | Mouse | 2D | manual | auto (Neuropixels probes/cells) | yes | - | DOI | https://github.com/cortex-lab/allenCCF |
| QUINT | 2019 | - | Linux, Windows | Mouse/Rat | 2D | with DeepSlice or QuickNII | semi-auto (cells (ilastiks)) | with MeshView | - | DOI | https://www.ebrains.eu/tools/quint-workflow |
| MIRACL | 2019 | Python | Linux | Mouse | 3D | auto | auto (cells (in ImageJ)) | yes | integration of cleared tissue with MRI data | DOI | https://miracl.readthedocs.io/en/latest/ |
| AMaSiNe | 2020 | Matlab | Windows | Mouse | 2D | auto | auto (cells) | yes | - | DOI | https://github.com/vsnnlab/AMaSiNe |
| BrainGlobe | 2020 -22 | Python | Linux, MacOS, Windows | Mouse/Rat/Zebrafish / (via bg_atlasapi) | 3D | auto (brainreg) | auto (cells/axonal projections/probes (CellFinder)) | yes (brainrender) | - | BG Atlas API<br>brainrender<br>CellFinder<br>brainreg/brainreg -segment | https://brainglobe.info/index.html |
| mBrainAligner | 2021 | C++ & Python | Linux, Windows | Mouse | 3D | auto (with curation) | no | no | comparing 3D brain imaged with different methods | DOI | https://github.com/Vaa3D/vaa3d_tools/tree/master/hackathon/mBrainAligner |
| SMART | 2022 | R | Linux, MacOS, Windows | Mouse | 3D | auto (with 'choice game') | yes (WholeBrain) | yes (WholeBrain) | extension of WholeBrain | DOI | https://mjin1812.github.io/SMART/ |
| SHARCQ | 2022 | Matlab | Linux, MacOS, Windows | Mouse | 2D | manual (SHARP-track) | external (ImageJ/Adobe) | yes | extension of SHARP-track | DOI | https://github.com/wildrootlab/SHARCQ |
| AMBIA | 2023 | Python | Linux, MacOS, Windows | Mouse | 2D | auto | yes (cells) | yes | - | DOI | https://github.com/mrymsadeghi/AMBIA |
| DeepSlice | 2023 | Python | Linux, MacOS, Windows | Mouse | 2D/3D | auto | no | no | integrated in QUINT workflow | DOI | https://github.com/PolarBean/DeepSlice |
| HERBS | 2023 | Python | Linux, MacOS, Windows | Mouse/Rat | 2D | manual | yes (probes/injection volumes) | yes (TRACER) | probe tract planning | DOI | https://github.com/Whitlock-Group/HERBS |
| ABBA | 2024 | Java | Linux, MacOS, Windows | Mouse/Rat/Zebrafish / (via bg_atlasapi) | 2D/3D | semi-auto | with BraiAn | with BraiAn | - | DOI | https://github.com/BIOP/ijp-imagetoatlas |
| Bell Jar | 2025 | Java | Linux, MacOS, Windows | Mouse | 2D | semi-auto | yes (cells) | no | - | DOI | https://github.com/asoronow/belljar |
| DMC-BrainMap | 2025 | Python | Linux, MacOS, Windows | Mouse/Rat/Zebrafish / (via bg_atlasapi) | 2D | manual | yes (cells/projections/injection volumes/Neurpixels probes (1.0)/optic fibers) | yes | - | - | https://github.com/hejDMC/napari-dmc-brainmap |
